# Data Denoising with transfer learning in single-cell transcriptomics

**DOI:** 10.1101/457879

**Authors:** Jingshu Wang, Divyansh Agarwal, Mo Huang, Gang Hu, Zilu Zhou, Chengzhong Ye, Nancy R. Zhang

## Abstract

Single-cell RNA sequencing (scRNA-seq) data is noisy and sparse. Here, we show that transfer learning across datasets remarkably improves data quality. By coupling a deep autoencoder with a Bayesian model, SAVER-X extracts transferable gene-gene relationships across data from different labs, varying conditions, and divergent species to denoise target new datasets.

---

In scRNA-seq studies, technical noise blurs precise distinctions between cell states, and genes with low expression cannot be accurately quantified. Existing methods^1–6^ to denoise scRNA-seq data often underperform when sequencing depth is low or when the cell type of interest is rare, and also ignore datasets in public domain, which may contain relevant information to aid denoising. Ensuing the mouse cell atlases^7,8^, we will soon have detailed atlases for each anatomic organ in the human body^9^. Publicly available scRNA-seq datasets contain information about cell types and gene signatures that is relevant to newly generated data. Yet, it is unclear how to borrow information across platforms, subjects and tissues. Moreover, such transfer learning must not introduce bias or force the new data to lose its distinctive features.

Here, we describe a denoising method, <u>S</u>ingle-cell <u>A</u>nalysis <u>v</u>ia <u>E</u>xpression <u>R</u>ecovery harnessing e<u>X</u>ternal data (SAVER-X), which couples a Bayesian hierarchical model to a pretrainable deep autoencoder^10^. Although neural networks have formed the basis of other single cell methods^4,6^, existing tools act solely on the data at hand. Moreover, extensive benchmarking^11^ and examples herein (Figure 3d, Figure S2) highlight that most methods except SAVER^1^, the precursor to SAVER-X, produce biased estimates of the true gene expression and introduce spurious gene-gene correlations. SAVER-X builds upon the core model from SAVER, combining it with the autoencoder backend with a two-stage training regime to utilize public data resources.

The statistical framework underlying SAVER-X is explicated in Methods. Briefly, let *Y* denote the new scRNA-seq count matrix to be denoised (“target data”). SAVER-X decomposes the variation in *Y* into: (i) a predictable structured component (*Λ*) which explains the shared variation across genes, (ii) unpredictable cell-level fluctuations that are independent across genes, and (iii) technical noise. SAVER-X estimates the unobserved true gene expression, *X*, centered at *Λ* with independent gene-specific dispersions:

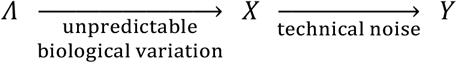

Λ is learned through an autoencoder (Figure 1b, item A), whose weights are first pretrained on cells from the same tissue or of similar type, extracted from public repositories (“pretraining data”; Figure 1a). The weights are then updated to fit the target data. This two-stage training regime allows adaptive retention of transferable features. Many core cell types and essential pathways are shared between human and mouse^9,12^. To allow cross-species learning, the autoencoder in SAVER-X includes a shared network between human and mouse (Figure S1). Additionally, SAVER-X employs cross-validation-based gene filtering and Bayesian shrinkage to preserve expression patterns that are unique to the target dataset (Figure 1b, items B and C). Cross-validation identifies genes poorly fit by the autoencoder, whose predictions are replaced by their target data mean. Bayesian shrinkage computes a weighted average of the predicted values 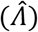 and the observed data (*Y*) to get the final denoised value 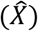.

**Figure 1.**
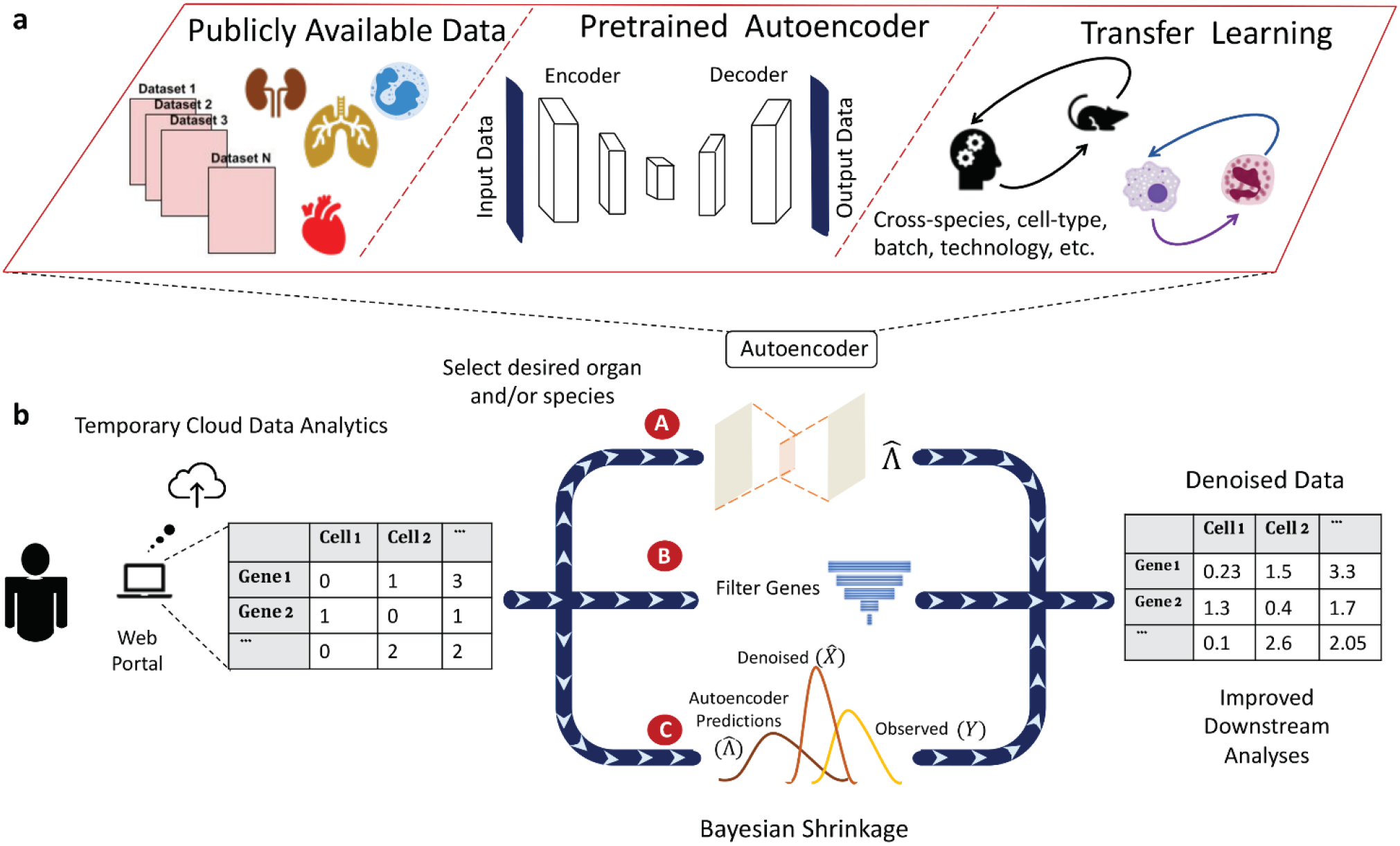
Outline of the SAVER-X transfer learning framework. **a**) The autoencoder pretraining step: for each species/organ/cell-type, public datasets are collected and combined to generate pretrained weights. **b**) Workflow of SAVER-X. Users can use SAVER-X web portal or run SAVER-X offline. For a target data with an UMI count matrix, SAVER-X trains the target data with autoencoder w/o. a chosen pretraining model (item A), then Filters unpredictable genes using cross-validation (item B) and estimates the final denoised values with empirical Bayes shrinkage (item C).

We first explored the benefits and limits of transfer learning via SAVER-X on a diverse testbed of cells that constitute the immune system. Despite being implicated in virtually every disease, tissue-infiltrating immune cells in scRNA-seq data are scarce in the absence of flow sorting. In such cases, denoising becomes especially challenging without the aid of external data ^13^. We examined whether SAVER-X pretrained on data from the Human Cell Atlas (HCA) project ^9^ (500,000 immunocytes from umbilical cord blood and bone marrow) and 10X Genomics ^14^ (200,000 Peripheral Blood Mononuclear Cells; PBMCs) could meaningfully improve the data quality of immune cells from healthy and disease tissue. Concomitantly, we benchmarked SAVER-X against existing denoising methods on a set of purified cells from 9 non-overlapping immune cell types ^14^.

Reliable identification of T-cell subtypes is crucial in characterizing a tissue’s immune environment, yet T-cell subtypes are often conflated in the raw scRNA-seq data (Figure 2a). We created a test dataset by randomly selecting 100 cells for each cell type, and found that SAVER-X pretrained on HCA not only markedly heightens the segregation among T-cell subtypes, but also improves the Adjusted Rand Index (ARI) over other methods (Figure S3). Encouraged by the transference of cell type-specific transcriptional signatures between datasets that contain similar cell types, we observed that the benefits of transfer learning manifest more clearly with decrease in cell number or sequencing depth in the target data (Figure S3). In the extreme case, even cells with a coverage of merely 60 total UMIs, typically discarded in current pipelines, can be rescued by transfer learning to reveal useful information.

**Figure 2.**
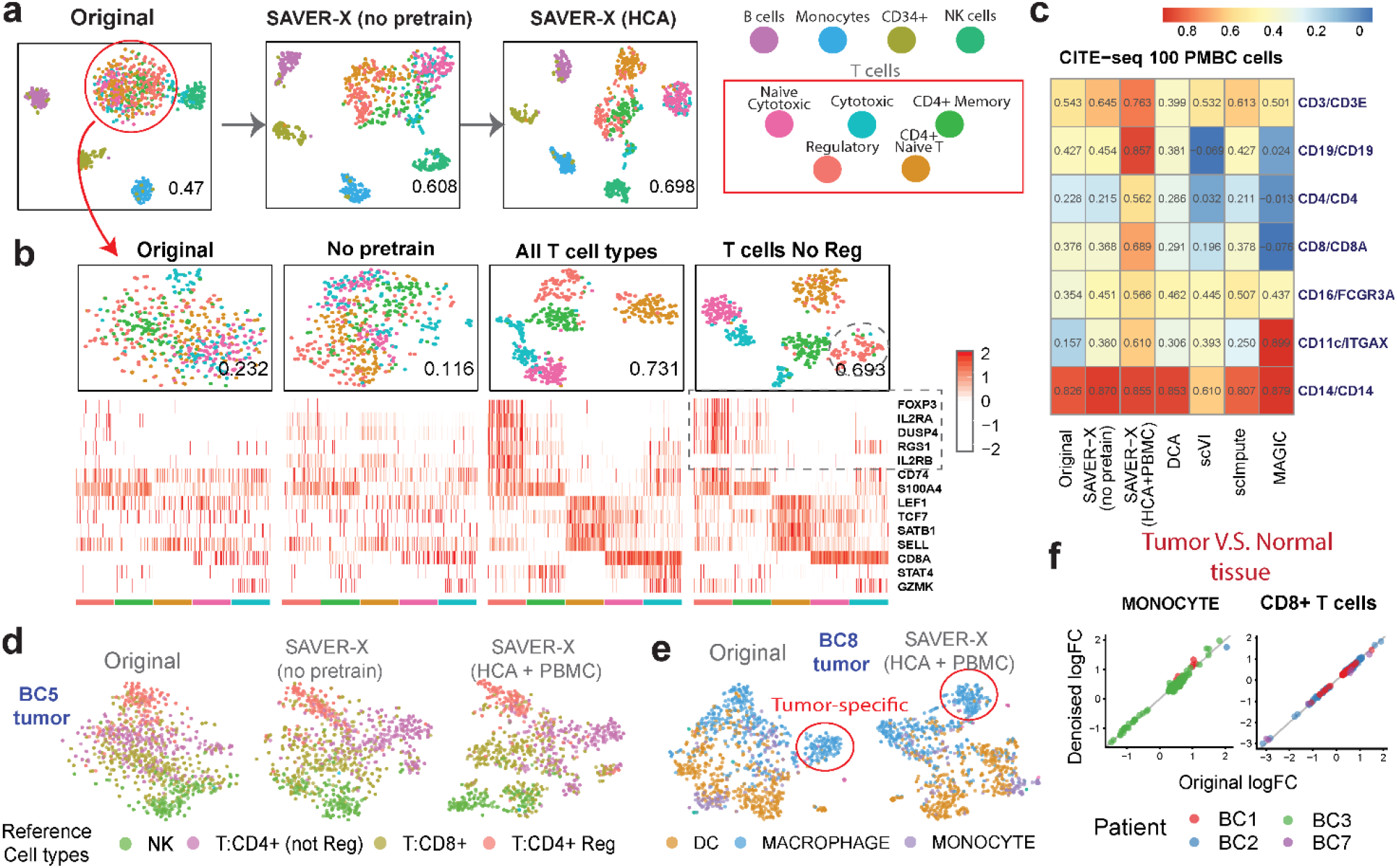
SAVER-X denoising of human immune cells. **a**) t-SNE plots of 900 immune cells, colored by known cell-type labels. The number at the right-bottom corner of each plot is the adjusted rand index (ARI). **b**) t-SNE plots of the T cells with different models (top) and the heatmap of the denoised gene log-scaled relative expressions for a set of known marker genes (bottom). **c**) Pearson correlations calculated between proteins and corresponding mRNA levels in the CITE-seq sub-sampled 100 PBMC cells after the RNA expressions are denoised using different methods. (**d**) T-SNE plots of the NK and T cells (1080 cells) for the BC5 tumor and (**e**) myeloid cells (1046 cells) for the BC8 tumor using SAVER-X with and without pretraining. Cells are colored with the cell types provided by the original paper. **f**) Log fold-change between paired tumor and normal tissues in 2 cell types and 4 patients. Patients BC1, BC2, BC3 and BC7 have 378, 53, 586 and 80 cells monocytes, and 4215, 1510, 58 and 401 CD8+ T cells respectively. The X-axis shows the log fold-change computed using the original data, and the Y-axis denotes that computed from the denoised data with normal immune cells pretraining. Each dot corresponds to a cell-type specific differentially expressed gene identified using the original data of each patient.

To appreciate the limits of transfer learning, we evaluated the relationship between denoising accuracy and cell type similarity between the pretraining and target datasets. Can transfer learning effectively denoise cellular states absent in the pretraining data? Consider the purified T-cells analyzed above. When SAVER-X is pretrained on all T-cell subtypes, clustering and expression quantification of marker genes improve substantially (Figure 2b). However, even when a cell type (CD4+ regulatory T-cells, T_regs_) is completely absent in pretraining, SAVER-X improves identification and marker gene quantification for this “new” cell type. Relatedly, to determine whether enrichment of a cell type in the pretraining data improves denoising accuracy in the target data, we pretrained SAVER-X on T-cells enriched for T_regs_, which did not produce any appreciable difference (Figure S4), Thus, SAVER-X does not require a perfect match in cell type composition between the pretraining and target data, and, importantly, can improve the quantification of new cell types that are absent in the pretraining data.

As an auxiliary measure, we also used CITE-seq ^15^ to examine the gene-protein correlations of key immune markers. Correlations between protein abundances and RNA expression of their cognate genes were found to be strikingly low in CITE-seq^15^. We discovered that for PBMC/CBMC CITE-seq data, the denoised expression estimates from SAVER-X (pretrained on HCA and PBMC 10X Genomics) have patently higher correlations with their protein product. Compared with other methods, SAVER-X consistently improved correlations across *all* markers when the target dataset contained 100 and 1000 cells (Figure 2c, S5). For larger data with 8000 cells, however, pretraining did not provide an apparent benefit (Figure S5).

Next, we probed whether SAVER-X can effectively learn from healthy immune cells to denoise immune cells sequenced from primary breast carcinoma samples^16^. Compared with the no-pretraining model, pretraining on immune cells from healthy tissue (HCA and PBMC 10X Genomics) allowed us to better characterize the tumor-infiltrating immune cell types in multiple subjects (Figure 2d, Figure S6). Meanwhile, a tumor-associated immune cell subpopulation remained identifiable after transfer learning. In particular, SAVER-X preserved the elevated immunoglobulin production signature in this disease-specific cell state (Figure 2e, Figure S7). This population was absent in the normal tissue, and we validated its immunophenotype by the presence of markers such as *LYZ*. Cell-type specific gene expression differences between paired tumor and normal tissues were also preserved in all patients with paired tissues and for both two cell-types vital in immune-surveillance (Figure 2f). These results highlight that SAVER-X can effectively harness public immune cell data from healthy conditions to denoise immune cells from disease conditions, while preserving disease-specific signatures.

Finally, we considered cross-species transfer learning using scRNA-seq data from cells in the developing ventral midbrain of both mouse and human^12^. We sampled 10% of the reads in the human data^12^, reducing it to a median-per-cell coverage of 452 UMIs, and utilized the original data as reference to gauge denoising accuracy. We split the human cells randomly into two groups, down-sampled the reads of one group and reserved the other group for pretraining (Figure S8a). SAVER-X pretrained on the matched mouse brain cells led to a noticeable improvement, compared to no-pretraining, in for human cell classification (Figure 3b). Pretraining on both human and mouse cells further improved the denoising accuracy compared to pretraining on human cells alone (Figure S8bd). Moreover, pretraining SAVER-X on cells from regions other than the ventral midbrain^7^ was beneficial, and so was pretraining on three human non-UMI datasets ^17–19^ together with mouse cells (Figure S8b). These experiments demonstrate the merit of transfer learning across species in general and practical settings where the anatomical regions and experimental protocols might differ between the pretraining and target data.

**Figure 3:**
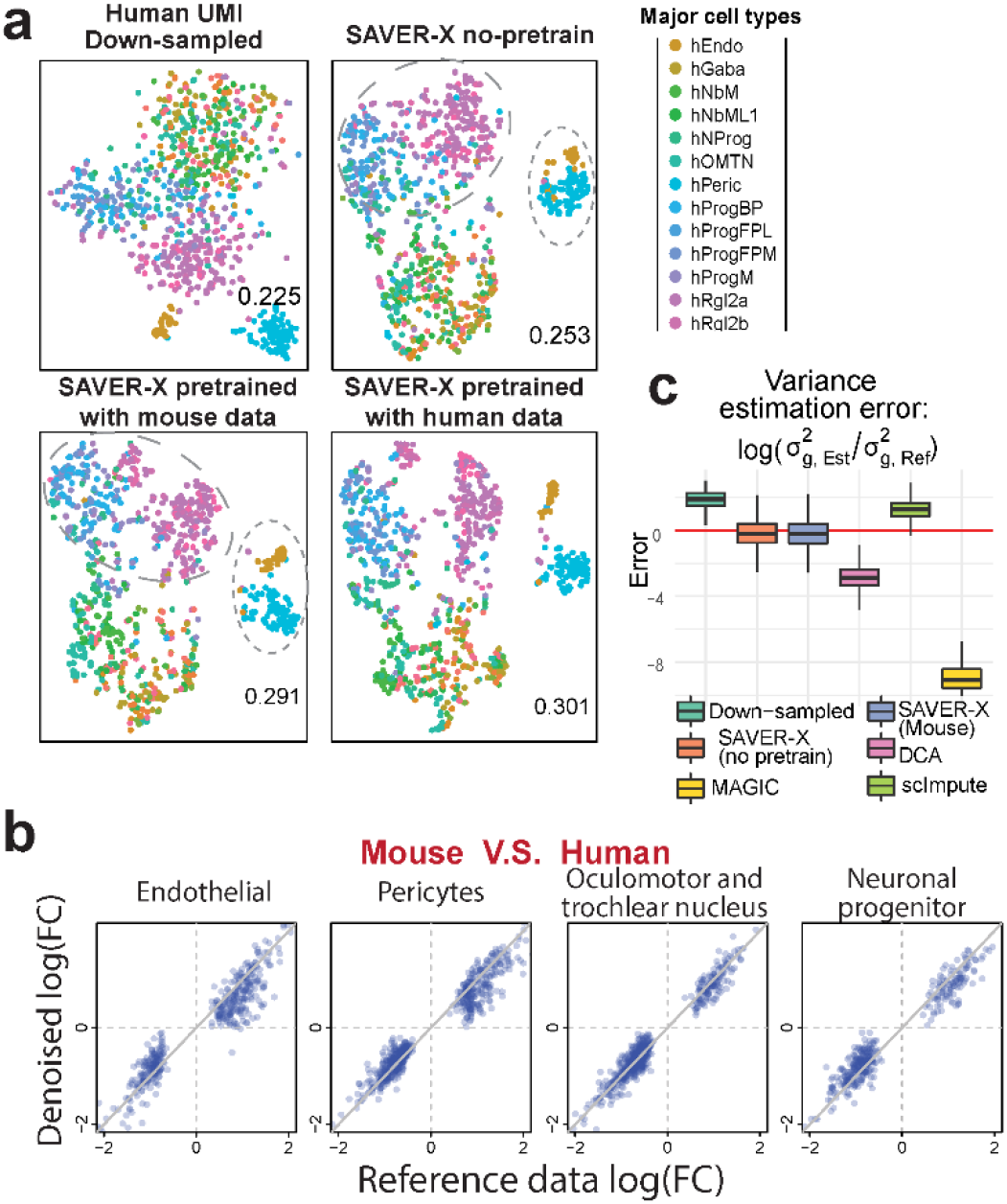
Mouse to human transfer learning within the developing ventral midbrain. **a**) t-SNE plots for the down-sampled data (n=1000 cells) and SAVER-X with different pretraining models. Reference cell labels are obtained from the original paper. The ARI displayed at the bottom-right corner of each plot is computed against the original labels. **b**) Log fold-change between human and mouse data in 4 major cell types. The X-axis shows the log fold-change computed using the original human data, and the Y-axis denotes the changes computed using the denoised down-sampled human data, wherein the denoising is done by SAVER-X pretrained with mouse cells. Each dot corresponds to a differentially expressed gene between human and mouse in that cell type. **c**) Box plots across all 20560 genes (showing the median, two hinges and two whiskers, outliers omitted) of the log ratios of the estimated variances compared with the reference data variances.

We then scrutinized whether a model pretrained on mouse data biases the estimates for genes with human-specific expression. We computed the log fold-change in cell type-specific average expression between human and mouse, and identified genes differentially expressed between the two species for four cell types. Denoising the down-sampled human data with SAVER-X pretrained on mouse cells preserved the log fold-changes (Figure 3b). In contrast, simply relying on the autoencoder, without cross-validation and shrinkage, reduced the fold-change for some genes (Figure S8c). Unlike other methods, SAVER-X also preserved the variance of genes across cells (Figure 3c).

Taken together, our results demonstrate that SAVER-X’s framework can leverage existing data to improve the quality of new scRNA-seq datasets. At its core, SAVER-X trains a deep neural network across a range of study designs, and applies this model to new data to strengthen shared biological patterns. Transfer learning changes the approach to scRNA-seq data analysis from a process of study-specific statistical modeling to an automated process of cross-study data integration and information sharing.

## Acknowledgements

We wish to thank Hugh MacMullan IV, Vincent Conley and Shawn Zamechek from Wharton Computing’s Research & Analytics Team (https://research-it.wharton.upenn.edu/) for their valuable assistance in implementing the code in a scalable fashion, and integrating the scalable code solution into a backend service for our website. We would also like to thank the National Institute of Health for the award 5R01-HG006137 (for D.A., Z.Z., and N.Z.), the National Science Foundation for the award DMS-1562665 (to J.W., N.Z.), the Blavatnik Family Foundation’s Graduate Student Fellowship awarded to D.A., NSF Graduate Fellowship DGE-1321851 awarded to M.H., and the Natural Science Foundation of Tianjin for grant 18JCYBJC24900 to G.H.. This work used the Extreme Science and Engineering Discovery Environment (XSEDE), which is supported by National Science Foundation grant number ACI-1548562. Specifically, it used the Bridges system, which is supported by NSF award number ACI-1445606, at the Pittsburgh Supercomputing Center (PSC).

## Author Contributions

J.W. and N.Z. conceptualized the study, designed the model and planned the case studies. J.W. developed the algorithm, implemented the SAVER-X software and led the data analysis. D.A. constructed the SAVER-X website and helped with data analysis. M.H. performed benchmarking with other methods and helped with algorithm development. G.H. helped with algorithm development and model design. Z.Z. tested SAVER-X website and software, and helped with data analysis. C.Y. conducted the analysis of CITE-seq data. J.W., D.A. and N.Z. wrote the paper with feedback from M.H. and Z.Z.

## Competing Financial Interests Statement

The authors declare no competing interests

## Online Methods

### Statistical model of SAVER-X

As stated in the main text, SAVER-X uses a statistical hierarchical model to decompose the randomness of observed UMI counts into three parts. Assume that the observed data is *Y* = (*Y_gc_*)_*G*×*C*_ where *g* represents a gene and *c* represents a cell. Also, assume that the true relative gene expression is *X* = (*X_gc_*)_*G*×*C*_ where *X_gc_* is the proportion of RNA copies of gene *g* in cell *c*. First, for the technical noise of UMI counts, it was shown ^20,21^ that a Poisson model,

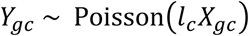

where *l_c_* = Σ*_g_ Y_gc_* is the library size of cell *c*, has substantial empirical and logical support. Thus, both SAVER and SAVER-X have adopted this model for the technical noise and only support denoising scRNA-seq data with UMI. Next, to quantify the biological variation, we assume that the true gene expression *X_gc_* is derived by adding independent fluctuations for each gene to an underlying gene-gene correlated component *Λ_gc_*, as follows:

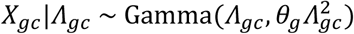

which is a Gamma distribution with mean *Λ_gc_* and variance 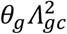. *Λ_gc_* can be interpreted as the portion of gene *g*’s expression that is predictable by other genes. Since, at the level of single cells, gene expression can be idiosyncratic and unpredictable, at least given our current knowledge, we let the true expression *X_gc_* to deviate from the correlated component *Λ_gc_* to better recover single-cell level expression stochasticity. This is a critical difference between our framework and other denoising frameworks.

### Pretraining and prediction with autoencoder

The autoencoder in SAVER-X is used to estimate *Λ* = (*Λ_gc_*)_*G*×*C*_. It can be trained exclusively on the target data set, or first pretrained on existing data sets and then on the target data set. To allow transfer learning across species, the autoencoder used by SAVER-X has three subnetworks, as shown in Figure S1, with one subnetwork taking human genes as input, one subnetwork taking mouse genes as input, and one subnetwork taking shared human-mouse homologous genes as input. 21183 and 21122 genes are used for human and mouse (Supplementary Data, Supplementary Note), respectively, as input and output nodes of the autoencoder. By current annotations using the getLDS() function in the bioMaRt R package, 15494 genes out of these nodes have homologs shared between the two species (Supplementary Data, Supplementary Note). For each sub-network, the number of nodes in the encoding and decoding layers are, successively, 128, 64, 32, 64, and 128. We find in practice that the results are relatively robust to the chosen number of layers and nodes.

For a given tissue or cell-type, and for a given (set of) specie(s), publicly available scRNA-seq data of the given category are mixed and fed into the autoencoder to pretrain a model. If only human data is available, only the human and shared sub-network weights are updated. Similarly, if only mouse data are available, only the mouse and shared sub-network weights are updated. Although SAVER-X can only denoise UMI counts, it can use both UMI and non-UMI datasets for pretraining. To adjust for the differences between data generated using non-UMI- and UMI-based technologies, an indicator node at the input layer feeds into each sub-network. For UMI datasets, the input expression levels are normalized by library size, re-scaled and log-transformed using formula: 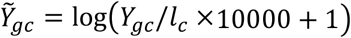 with 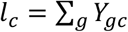. For non-UMI datasets, TPM for each cell *c* and gene *g* are denoted as *Y_cg_*, and then the *Y_cg_* are transformed using the same formula as that for UMI. We use the negative log-likelihood function of the data as the loss function of our autoencoder. For UMI counts, based on our hierarchical model, *Y*|*Λ* follows a Negative Binomial distribution. Thus, for each cell c in one dataset, the loss function is:

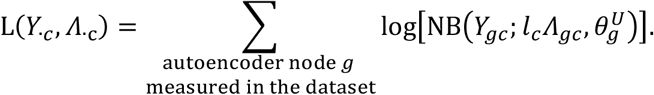

On the other hand, for the non-UMI TPM data, *Y*|*Λ* is assumed to approximately follow a zero inflated Negative Binomial distribution (although TPM is not integer-valued, the likelihood function can still be computed) and the loss is:

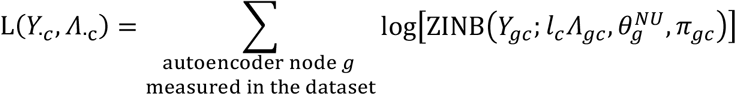

Here NB(*x*; *μ*, *θ*) and ZINB(*x*; *μ*, *θ*, *π*) are the densities of Negative Binomial and zero-inflated Negative Binomial distribution (see Supplementary Note). A separate gene-specific dispersion parameter 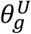 and 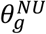 is dedicated for UMI and non-UMI input, respectively. For non-UMI data, the gene- and cell-specific zero inflation parameter is defined as

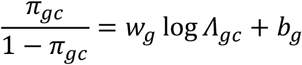

Where *w_g_* and *b_g_* are gene-specific unknown coefficients. SAVER-X only transfers the gene-gene relationship weights, and does not transfer the over-dispersion nor zero inflation parameters. Our implementation of the autoencoder utilizes the library functions of DCA ^4^.

### Cross-validation and filtering for unpredictable genes

When training the autoencoder on a target UMI dataset to get the prediction matrix 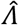, cross-validation is applied to filter out genes that cannot be predicted accurately by the autoencoder. Specifically, the target data is randomly split into held-in and held-out cell sets, the autoencoder is trained on the held-in set and then used to make predictions on the held-out set. We compare the performance of the autoencoder with a completely null model where the gene expression in every cell is predicted by their means across cells. For a specific gene *g*, let the held-in sample mean for the library-size normalized counts be *μ_g_*. Then a gene is unpredictable if the Poisson deviance of the autoencoder is larger than that of the null model, equivalently:

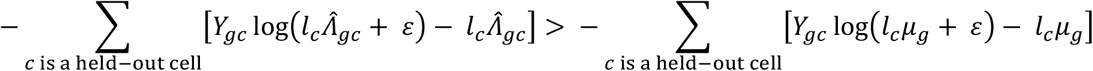

where *ε* = 10^−10^ to avoid taking the log of zeros. For the unpredictable genes that are identified by cross-validation, their predictions are replaced with the null model prediction, which are the sample means of library size normalized UMI counts for every gene.

### Empirical Bayes Shrinkage

After estimating 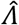, the final denoised matrix 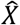 of SAVER-X is obtained by empirical Bayes shrinkage based on the hierarchical model:

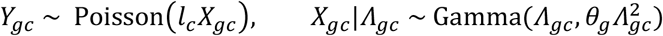

We replace the unknown *Λ_gc_* with the estimated 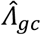 and estimate *θ_g_* for each gene by maximum likelihood. The final denoised value 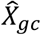 is the posterior mean of *X_gc_*, which is an inverse-variance weighted average:

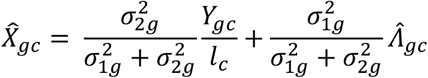

where 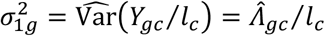 and 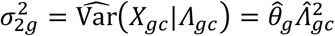. SAVER-X also outputs the posterior variance of each *X_gc_*.

### Data denoising using other bench-marking methods

MAGIC ^3^ was performed using the R version 1.3.0 on the square root transformed mean library-size normalized expression. scImpute ^2^ version 0.0.9 was performed on the unnormalized expression values. The tuning parameter Kcluster is set to 9 for the PBMC data ^14^, set to 11 for the CITE-seq data, and set to 13 for the human midbrain data. DCA ^4^ version 0.2.2 was performed on the unnormalized expression values and the library-size normalized expression output was used for downstream analysis. scVI ^6^ version 0.2.4 was performed with n_epochs = 400, traing_size = 0.9, frequency = 5 and lr = 1e-3 and all genes being used.

### Generating down-sampled datasets

For an observed UMI count data matrix, we down-sample the reads to obtain a data set of the same gene and cell numbers but with lower quality. For cell *c* and gene *g*, we treat the original count as the true expression *X_gc_* and the down-sampled value *Y_gc_* is generated following the Poisson-alpha technical noise model in Wang et. al. ^20^ by independently drawing from a Poisson distribution with *Y_gc_*~Poisson(*l_c_X_gc_*) where *l_c_* is a cell-specific efficiency loss. To mimic variation in efficiency across cells, we sampled *l_c_* as follows:

1. 10% efficiency: *l_c_*~Gamma(10,100), used on the mouse midbrain data ^12^
2. 5% efficiency: *l_c_*~Gamma(5,100), used on the 10X PBMC data ^14^

### t-SNE visualization and cell clustering

We used Seurat version 2.0 to perform cell clustering and t-SNE visualization according to the preprocessing workflow detailed at (https://satijalab.org/seurat/pbmc3k_tutorial.html). For all analyses, we set the number of principal components (PC) to 15, though we find our comparison results robust to a range of PC from 10 to 20. For cell clustering using Seurat, resolution is set to be 1.6, 1.2, 0.8 and 0.8 for each of the four experiments (90 cells, 900 cells, 9000 cells and 9000 cells with down-sampled reads) of the PBMC data (Figure 2ab, Figure S2) and kept the same on all the methods compared. The resolution is set to 1 for the cell clustering of the PBMC T cells (Figure 2c, Figure S4) and the midbrain data ^12^. The adjusted Rand Index (ARI) is computed using R package mclust to evaluate the clustering performance compared with the known cell type labels. We exclude human midbrain cells that are labeled as either “unknown” or three rare types (“hMgl”, “hOPC”, “hSert”) in the original paper for both the t-SNE plot and ARI calculation.

Each t-SNE plot is generated after going through a fresh preprocessing workflow of the cells in the plot using Seurat. For some t-SNE plots, companion feature plots are shown for selected marker genes (Figure S7, Figure S7) using Seurat 2.0 function FeaturePlot().

### Analysis of the CITE-seq data ^15^ and Breast Cancer data ^16^

The CITE-seq data were pre-processed as described in the original paper. Cells with more than 90% UMI counts from human genes were retained. The mouse genes and zero expression genes were removed. To study the situation where the information available from the target data is limited, we created two subsampled data with 1,000 and 100 cells from the PBMC and CBMC data, respectively, along with the original datasets (~8,000 cells for each). The mRNA data were then denoised using SAVER-X without pretraining, SAVER-X pretrained with immune cells (HCA + PBMC) and other benchmarking methods. mRNA data before and after denoising were log-normalized using Seurat 2.0. The ADT (antibody-derived tag) protein measurements were normalized by the authors using centered-log-ratio (CLR) transformation. Pearson correlation were then calculated between the transformed and normalized proteins and mRNA.

The breast cancer data is denoised using SAVER-X pretrained immune cell model for each patient and each tissue separately. Cell type labels of the original paper are used as reference cell types subjecting to estimation inaccuracies of the original paper.

### Differential expression analysis

There are two sets of differential expression (DE) analyses, DE between the tumor and normal tissue for each patient where both tissues are measured (Figure 2f), and DE between human and mouse of the developmental midbrain (Figure 3c, Figure S8c). For both, the DE analysis is done for each cell type separately using the original UMI counts and cell types provided by the original paper. The datasets are preprocessed in Seurat 2.0 following standard workflow and the DE genes are also obtained using Seurat 2.0, where the Wilcoxon rank sum test is used. P-value adjustment is performed using Bonferroni correction based on the total number of genes in the dataset. A gene is selected as differentially expressed if its adjusted p-value is ≤ 0.05 and the absolute log fold change is ≥ 0.25.

### Variance calculation and comparison

For the human developmental midbrain data, we compared the variances of each gene estimated using different denoising methods with the sample variances directly calculated from the reference data, which are considered as true values in our down-sampling experiment. Different denoising methods are applied to the down-sampled human midbrain dataset. Specifically, we calculate and compared the variances of the relative gene expression across cells. For other denoising methods, the variances are estimated as the sample variance of the denoised and normalized data. Since SAVER-X outputs a posterior distribution:

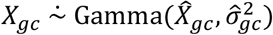

where 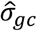 is the estimated posterior standard deviation, we estimate the variance of *X_gc_* across cells as the sample variance of 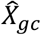 plus the average of 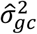 across cells. The differences between the estimated variances of each gene and the true variances calculated from the reference data are shown in Figure 3d. The down-sampled raw data has variances biased upwards because of the technical noise while DCA and MAGIC have variances biased downwards due to over-smoothing.

### Data Availability

The Human Cell Atlas (HCA) dataset was downloaded from the HCA data portal (https://preview.data.humancellatlas.org/) and the PBMC data ^14^ was downloaded from the 10X website (https://support.10xgenomics.com/single-cell-gene-expression/datasets, Table S2). The breast cancer data ^16^ was downloaded from GEO (GSE114725). The developing midbrain data ^12^ was downloaded from GEO (GSE76381). For the other mouse developing brain datasets in Figure 3, we include cells from neonatal and fetal brain tissues in the Mouse Cell Atlas ^7^ data (GSE108097). For the non-UMI human developing brain datasets in Figure S8, we include three: GSE75140 ^18^, GSE104276 ^19^ and SRP041736 ^17^. No gene or cell filtering is done on the original dataset.

A complete list of the pretraining datasets used for pretraining the models on the SAVER-X website is provided in Table S2.

### Software Availability

SAVER-X is publicly available at http://singlecell.wharton.upenn.edu/saver-x/, where users can currently upload their data for cloud computing and choose from models pre-trained on 31 mouse tissues and human immune cells. Models jointly pretrained on cells from both species are also available for brain and pancreatic tissues. The R package and source code of SAVER-X is also released at https://github.com/jingshuw/SAVERX.

## Notes

#### Summary of Updates

Title, main text and supplementary materials changed

