## supplementary text, tables and figures for "Data Denoising with transfer learning in single-cell transcriptomics"

**Supplementary note 1**: Consensus gene mapping of the datasets for autoencoder pretraining

In order to pretrain multiple datasets jointly at the same time and to enable transferring similar gene-gene relationships to the target data, we need to first map the gene names in each dataset to a consensus list of gene names. Currently, some genes do not have unique names and in SAVER-X, we create a mapping function to map possible gene names to a list of standard names to our best knowledge.

For human, such a standard name list contains all the HGNC symbols (complete HGNC dataset downloaded from https://www.genenames.org/cgi-bin/statistics) and Ensemble gene IDs for those genes without an HGNC symbol. For mouse, the consensus list contains all the MGI symbols (downloaded from http://www.informatics.jax.org/downloads/reports/index.html) and Ensemble gene IDs for those genes without MGI symbols. Ensemble gene IDs for both human and mouse were obtained using the useMart() function in the biomaRt R package. For the gene names in each dataset that are not exactly one of the names on the list, we checked for additional information on previous gene names, alias gene names and wiki gene names from the reference websites above and from the biomaRt R package. This provided us with the maximum number of genes that can be mapped to our consensus list.

Then, we also need to determine what genes we include as an input/output node in the SAVER-X autoencoder. For both species, a gene from the consensus list is selected as one autoencoder node if it appears in at least half of all our pre-training datasets. Currently, we have 21183 genes for human and 21122 genes for mouse. An autoencoder gene node is further determined to be shared in both species if both its mapping from human to mouse and vice-versa are unique using the getLDS() function in the bioMaRt R package. Currently, we have 15494 genes that are shared. Complete lists of the human, mouse and shared gene nodes are stored as datasets (data(human_nodes_ID), data(mouse_nodes_ID), data(shared_nodes_ID)) in the SAVERX R package.

When we pretrain or train the autoencoder for a dataset, if a gene node is not uniquely measured in the dataset, we mark it as not measured (missing). We set the input of the not measured genes as 0 and exclude these genes when calculating the loss when training the autoencoder. For a target dataset, the predicted values of the genes not passed to the autoencoder will be their cross-cell averages, which are the predictions of the null model. Thus, we can take all the genes to the Bayesian shrinkage step and our final denoised data matrix has the same size as the original input matrix.

**Supplementary note 2**: SAVER-X choice of autoencoder tuning parameters and optimization details

As a neural network, the autoencoder has a lot of tuning parameters. We have not tried all possibilities but have chosen the set of tuning parameters that currently gives satisfying empirical results.

We set 3 encoder layers and 3 decoder layers for each of the sub-neural network. A larger autoencoder will be generally easier for the gradient descent methods to find optimal solution, but it also takes more RAM and longer time. We choose the three hidden layers to have 128, 64 and 32 nodes respectively (Figure S1) to reach a balance. In practice, we the find the down-stream analysis results to be relatively robust to the choice of the number of nodes and layers. All layers are fully connected dense layers.

We follow most of the optimization settings same as the default settings of DCA [1]. We use the RELU activation function with batch normalization. We do not use dropout, as finding it not improving optimization. Early stopping is used with 10% of the cells randomly chosen as a validation dataset. "RMSprop" in Keras is used as the optimization method. The pretraining batch size is fixed as 100 cells. The batch size for training the target dataset is adaptively set as max(32, number of cells/50) or can be user-specified. The autoencoder weights are initialized randomly using the "glorot_uniform" algorithm in Keras when trained without a pretraining model.

**Supplementary note 3**: Density function forms for Negative Binomial, zero-inflated Negative Binomial and Gamma distributions

$$\mathrm{NB}(x;\mu,\theta) = \frac{\Gamma(x+\theta)}{\Gamma(\theta)\Gamma(x+1)}\left(\frac{\theta}{\theta+\mu}\right)^{\theta}\left(\frac{\mu}{\theta+\mu}\right)^{x}$$

$$\mathrm{ZINB}(x;\mu,\theta,\pi) = \pi\delta_0(x) + (1-\pi)\mathrm{NB}(x;\mu,\theta)$$

$$\mathrm{Gamma}(x;\alpha,\beta) = \frac{\beta^{\alpha}}{\Gamma(\alpha)}x^{\alpha-1}e^{-\beta x}$$

**Supplementary note 4:** In silico experiments on the PBMC T cell transfer learning

In order to understand the performance and robustness of transfer learning under different pretraining scenarios, we design a fully controlled experiment. The experiment uses PBMC data (both the purified cells and a mixed population of ~68,000 cells from a fresh donor) from Zheng et. al. [2]. Based on the original paper, the cell type labels of the purified cells are obtained by conventional trustable approaches while the cell types of the mixed cell population are estimated from the same single-cell data. Thus, the target dataset we use is a population of 500 T cells, consisting of 100 randomly selected purified cells from each of the five non-overlapping T cell types.

For the pretraining dataset, we consider 4 scenarios:

1. **All the non-T PBMC cells**: including 30,314 cells from the 4 non-T purified cell types and 15,109 cells from the mixed population that are labeled as non-T cells.
2. **T cells all types**: including randomly sampled 5,000 cells from each of the purified T cell population
3. **T cells but without regulatory T cells**: the second scenario but without the 5,000 CD4+/CD25+ Regulatory T Cells
4. **T cells with regulatory T cells enriched**: the second scenario and with all other purified CD4+/CD25+ regulatory T cells and 14,112 cells from the mixed population that are labeled as CD4+/CD25+ regulatory T cells.

Here the pretraining data and target data are sequenced using the same protocol, in the same lab and with similar sequencing depth, thus we focus on understanding what can be transferred when the cell type population is different between the target and pretraining dataset.

We choose the regulatory T cells as they are hard to be separated from the CD4+ memory T cells using scRNA-seq data. The last two scenarios can help use understand whether we need all cell types to be presented in the pretrained data and whether enriching a cell type in the pretraining dataset can help denoising that cell type in the target data. The results of this experiment (Figure 2c, Figure S4) show that transfer learning is quite robust to these different scenarios. Even in the first scenario pretraining can help, since the non-T immune cells shared some similar characteristics as the T cells (such as gene characteristics of the naïve state of immune cells).

**Supplementary note 5**: Comparison between SAVER-X without pre-training and the autoencoder-based denoising methods DCA [1] and scVI [3]

All three methods: SAVER-X, DCA and scVI are based on the autoencoder architecture. However, besides the key feature that SAVER-X transfers from external data, SAVER-X also differs from the other two approaches by employing cross validation and by using an empirical Bayes shrinkage step. The empirical Bayes shrinkage estimates the final denoised expression levels as weighted averages between the autoencoder outputs and the observed counts.

The empirical Bayes step is based on the assumption that the technical noise in UMI-based scRNA-seq counts follows a Poisson-alpha model, which is supported with extensive empirical experiments using the ERCC spike-in genes from 9 public datasets [4]. Through down-sampling experiments with the Poisson-alpha technical noise model on four public datasets [5–8], this final empirical Bayes step is found to be effective in bias reduction in the denoised matrix for SAVER-X. Furthermore, we show that empirical Bayes shrinkage effectively reduces bias even when coupled with DCA and scVI.

To produce the down-sampled data for each dataset, we first select high-quality cells and genes with high expression (details see Online Methods of Huang et al. [9]) from the original dataset to treat as the true expression $\lambda_{cg}$. Then we generated down-sampled datasets following the Poisson-alpha noise model, $x_{cg} \sim \mathrm{Poisson}(l_c \lambda_{cg})$, by randomly drawing from a Poisson distribution with mean parameter $l_c \lambda_{cg}$, where $l_c$ is the cell-specific efficiency loss. As in Huang et al., to mimic variation in efficiency across cells, we sampled $l_c$ as follows:

1. $10\%$ efficiency: $l_c \sim \mathrm{Gamma}(10, 100)$, used on Baron et al. [6], Chen et al. [7] and La Manno et al. [8]
2. $5\%$ efficiency: $l_c \sim \mathrm{Gamma}(10, 200)$, used on Zeisel et al. [10]

We calculate the gene-gene correlations in the denoised matrix for all gene. For each gene pair, we calculate the difference between their correlation in the denoised matrix and in the reference matrix (the original matrix before down-sampling). For SAVER-X and DCA/scVI with empirical Bayes shrinkage, we calculate post-denoising adjusted correlations, using both the estimates and posterior variances (see Online Methods of Huang et al. [9]). Figure S1 shows the density plots of these differences in each of the four datasets. A density plot that is shifted towards the positive direction reveals that the method introduces spurious relationships between genes and produces inflated correlation estimates. A density plot that is tightly concentrated around zero signifies that the method gives unbiased estimates of gene-gene correlations.

We compare SAVER-X without pretraining with DCA and scVI. For DCA, we use their default values for the tuning parameters. For scVI, we keep all genes in the data as inputs to their autoencoder to allow a fair comparison and set all other tuning parameters to their default values. First, note that the differences for SAVER-X are tightly concentrated around zero in all four data sets, thus demonstrating that SAVER-X does not spuriously inflate correlation. Furthermore, note that both DCA and scVI produce severely inflated correlations that are larger than the true gene-gene correlations. However, if we add a final empirical Bayes shrinkage step to either DCA or scVI, like the last step of SAVER-X, this bias in gene-gene correlations can be greatly reduced (density curves are more concentrated around 0).

This analysis shows that direct estimates obtained from autoencoders can be biased and should be treated with caution. With properly modeled technical noise, the empirical Bayes shrinkage step in SAVER-X can greatly reduce bias in the denoised data.

**Supplementary note 6:** Data integration after denoising with SAVER-X

Currently, SAVER-X cannot remove the batch effects in the target data. However, it can be combined with current data integration and alignment methods to remove batch effects after data denoising. We can achieve that by simply align the denoised data matrices using existing data integration/alignment methods.

We considered a scenario where we attempt to align cells from healthy human peripheral and cord blood tissues (PBMC and CBMC, respectively) [11]. We used SAVER-X immune cell pretraining model to denoise both the CBMC and PBMC datasets separately, each of which contains ~8,000 cells. As illustration (Figure S9a), we used a popular data integration pipeline, Seurat CCA version 2 [12], to align the raw data using the first 20 CCs. This allowed us to identify the major cell types in the blood and demonstrated a good concordance of these types among the two datasets. We used marker genes for each of the major cell types as described in the original paper [11] to ascribe cell type identity to the identified clusters. We then used the Seurat CCA pipeline to align the SAVER-X denoised PBMC and CBMC datasets also using 20 CCs and identifying the cell types based on the marker gene expression profiles (Figure S9b). We found that not only is it feasible to align denoised data using the existing pipeline, but that the alignment of the denoised datasets also retains many of the salient visualization features that emerge when the two raw datasets are aligned. We see that denoising improves the visualization of the B cell clusters and was helpful in separating CD4+ T cells from CD8+ T cells. Comparison of the top and bottom left panels in Figure S9 shows that the identified clusters, for instance, for NK cells, monocytes and T cells, clearly contain cells from both the datasets, suggesting that the denoising process does not bias the existing data alignment pipelines.

Altogether, this analysis component demonstrated that denoised data can be readily used in existing scRNA-seq data alignment tools.

**Table S1:** Testbed to illustrate the performance of SAVER-X and transfer learning.

| Pre-training data | Test Data | Transfer across… | Design | Validation metric |
|---|---|---|---|---|
| HCA (bone marrow, cord blood) | 10x website, purified data of 9 cell types | Lab, tissue | Reduce cell number, Subsampling of reads | Known cell type labels, stability of known marker genes |
| 10x PBMC T cells, all samples | 10x T cells, labeled samples | Samples | Reduce cell number, Subsampling of reads | |
| 10x PBMC + HCA | Azizi et.al. (2018) breast cancer tumor cells | Lab, tissue, technology, condition | No subsampling. | Cell type labels from the original paper, known marker gene enrichment |
| Mouse cells from La Manno et. al. (2016). | Human cells from La Manno et. al. (2016). | Species | Subsampling of reads | Cell type labels from the original paper, compare to original full data, preserving fold change |
| Mouse brain cells from "*Tabula Muris*" | | Species, anatomical regions, subject ages, labs | | |
| Human + mouse cells from La Manno et. al. (2016). | | Species | | |
| Human + mouse cells from multiple published datasets | | Species, anatomical regions, subject ages, labs, technology | | |

**Table S2**: Pre-training datasets for each pre-trained models on the SAVER-X web portal.

| Tissue / Cell type | | Datasets |
|---|---|---|
| Mouse Models | Bladder | GSE108097 [1] (GSM2889480) |
| | Bone Marrow | GSE108097 [1] (GSM2906396, GSM2906399 - GSM2906404) |
| | Adult Brain | GSE93374 (Campbell 2017 [2]), GSE75330 (Marques 2016 [3]), GSE74672 (Romanov 2017 [4]), GSE71585 (Tasic 2016 [5]), GSE60361 (Zeisel 2015 [6]), GSE59739 (Usoskin 2015 [7]), GSE87544 (Chen 2017 [8]), GSE108097 [1] (GSM2906405, GSM2906406) |
| | Developing Brain | GSE76381 (La Manno 2016 [9]), GSE108097 [1] (GSM2906415, GSM2906454, GSM2906455) |
| | Calvaria | GSE108097 [1] (GSM2906445, GSM2906446) |
| | Embryo | GSE57246 (Biase 2014 [10]), GSE45719 (Deng 2014 [11]), GSE53386 (Fan 2015 [12]) |
| | Mesenchyme | GSE108097 [1] (GSM2906412) |
| | Embryonic Stem cells | GSE65525 (Klein 2015 [13]), GSE108097 [1] (GSM2906413, GSM2906414, GSM2935549) |
| | Gonads | GSE108097 [1] (GSM2906416, GSM2906423) |
| | Heart | GSE108097 [1] (GSM2906447) |
| | Hematopoietic Stem cells | GSE76983 (Grun 2016 [14]) |
| | Intestine | GSE108097 [1] (GSM2906417, GSM2906467 - GSM2906470) |
| | Kidney | GSE107585 (Park 2018 [15]), GSE108097 [1] (GSM2906418, GSM2906419, GSM2906425, GSM2906426) |
| | Liver | GSE108097 [1] (GSM2906421, GSM2906427, GSM2906428) |
| | Lung | GSE108097 [1] (GSM2906422, GSM2906429 - GSM2906431) |
| | Mammary Gland | GSE108097 [1] (GSM2906432 - GSM2906442, GSM2889483, GSM2889484) |
| | Muscle | GSE108097 [1] (GSM2906444, GSM2906448, GSM2906449) |
| | Ovary | GSE108097 [1] (GSM2906456, GSM2906457) |
| | Pancreas | GSE84133 (Baron 2016 [16]), GSE108097 [1] (GSM2906458, GSM3004530, GSM3004531) |
| | Peripheral Blood | GSE108097 [1] (GSM2906459 - GSM2906464) |
| | Placenta | GSE108097 [1] (GSM2906465, GSM2906466) |
| | Prostate | GSE108097 [1] (GSM2906481, GSM2906482) |
| | Retina | GSE63472 (Macosko 2015 [17]), GSE81904 (Shekhar 2016 [18]) |
| | Rib | GSE108097 [1] (GSM2906450 - GSM2906452) |
| | Skin | GSE108097 [1] (GSM2906453) |
| | Spleen | GSE108097 [1] (GSM2906471) |
| | Stomach | GSE108097 [1] (GSM2906424, GSM2906472) |
| | Testis | GSE108097 [1] (GSM2906473, GSM2906474) |
| | Thymus | GSE108097 [1] (GSM2906475, GSM2906476) |
| | Trophoblast Stem cells | GSE108097 [1] (GSM2906477) |

| | | |
|---|---|---|
| | Uterus | GSE108097 [1] (GSM2906478, GSM2906479) |
| Joint Species (shared) Models | Adult Brain | **Human**: (Lake 2016 [19])<br>**Mouse**: GSE93374 (Campbell 2017), GSE75330 (Marques, 2016), GSE74672 (Romanov 2017), GSE71585 (Tasic 2016), GSE60361 (Zeisel 2015), GSE59739 (Usoskin 2015), GSE87544 (Chen 2017), GSE108097 (GSM2906405, GSM2906406) |
| | Developing Brain | **Human**: GSE75140 (Camp 2015 [20]), GSE76381 (La Manno 2016 [9]), SRP041736 (Pollen 2014 [21]), GSE104276 (Zhong 2018 [22])<br>**Mouse**: GSE76381 (La Manno 2016), GSE108097 (GSM2906415, GSM2906454, GSM2906455) |
| | Pancreas | **Human**: GSE84133 (Baron 2016 [16]), GSE85241 (Muraro 2016 [23]), E-MTAB-5061 (Segerstolpe 2016 [24]), GSE81608 (Xin 2016)<br>**Mouse**: GSE84133 (Baron 2016), GSE108097 (GSM2906458, GSM3004530, GSM3004531) |
| Human Models | Immune cells (both innate and adaptive) | HCA, 10X website (4k and 8k from a healthy donor, aggregate of t_3k and t_4k, fresh 68k PBMC, purified data for each cell type from Zheng 2017 [25]) |
| | T cells | T cells of the PBMC cells from the 10X website |

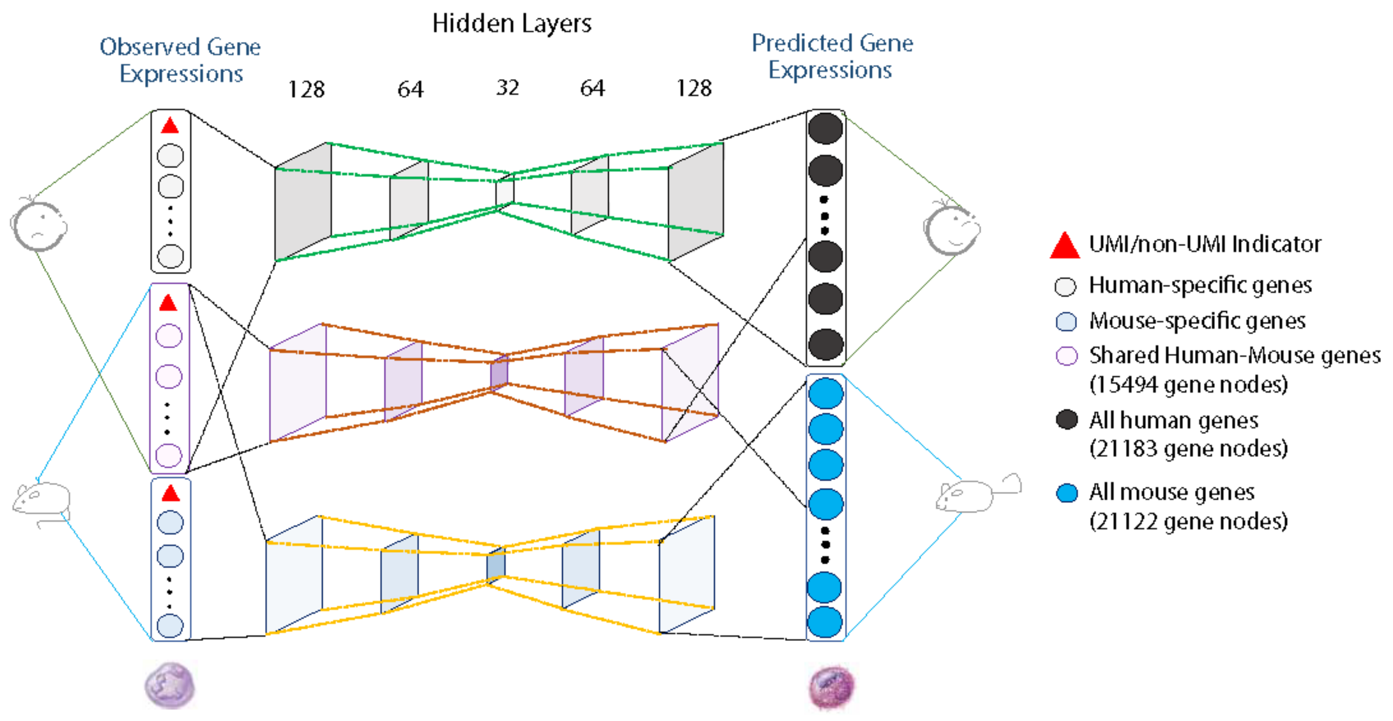


**Supplementary Figure 1**

Architecture of the autoencoder.

The autoencoder allows cells from both human and mouse, both with and without UMI, to be used for pre-training. For each species, there are ~20,000 input nodes accepting raw gene expression values; two-thirds of these are shared between mouse and humans by accounting for genes with homologs. See supplementary note 1 for more details.

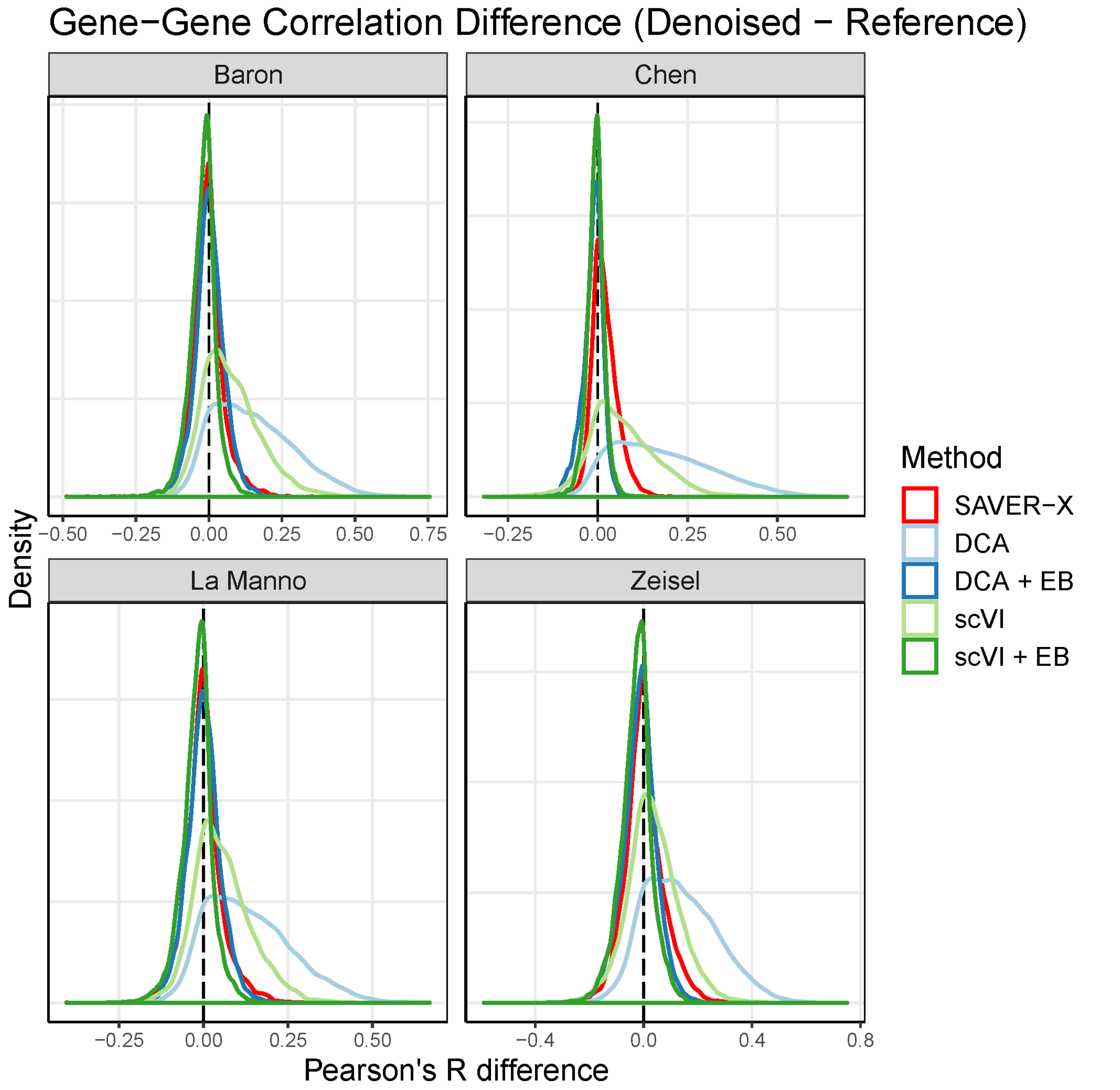


**Supplementary Figure 2**

Gene-gene correlation recovery.

Gene-gene Pearson correlations of all gene pairs are calculated in the denoised matrix and compared with the gene-gene correlations of the reference normalized count matrix. These are density plots of the difference between the denoised matrix and the reference across all gene-gene pairs for four different datasets. We compare five different methods: SAVER-X, DCA, scVI and the latter two with empirical Bayes shrinkage as in SAVER-X (DCA + EB, scVI + EB). These plots show that our final empirical Bayes shrinkage step in SAVER-X is essential to reduce bias in the denoised data regardless of the denoising method. See supplementary note 5 for more details.

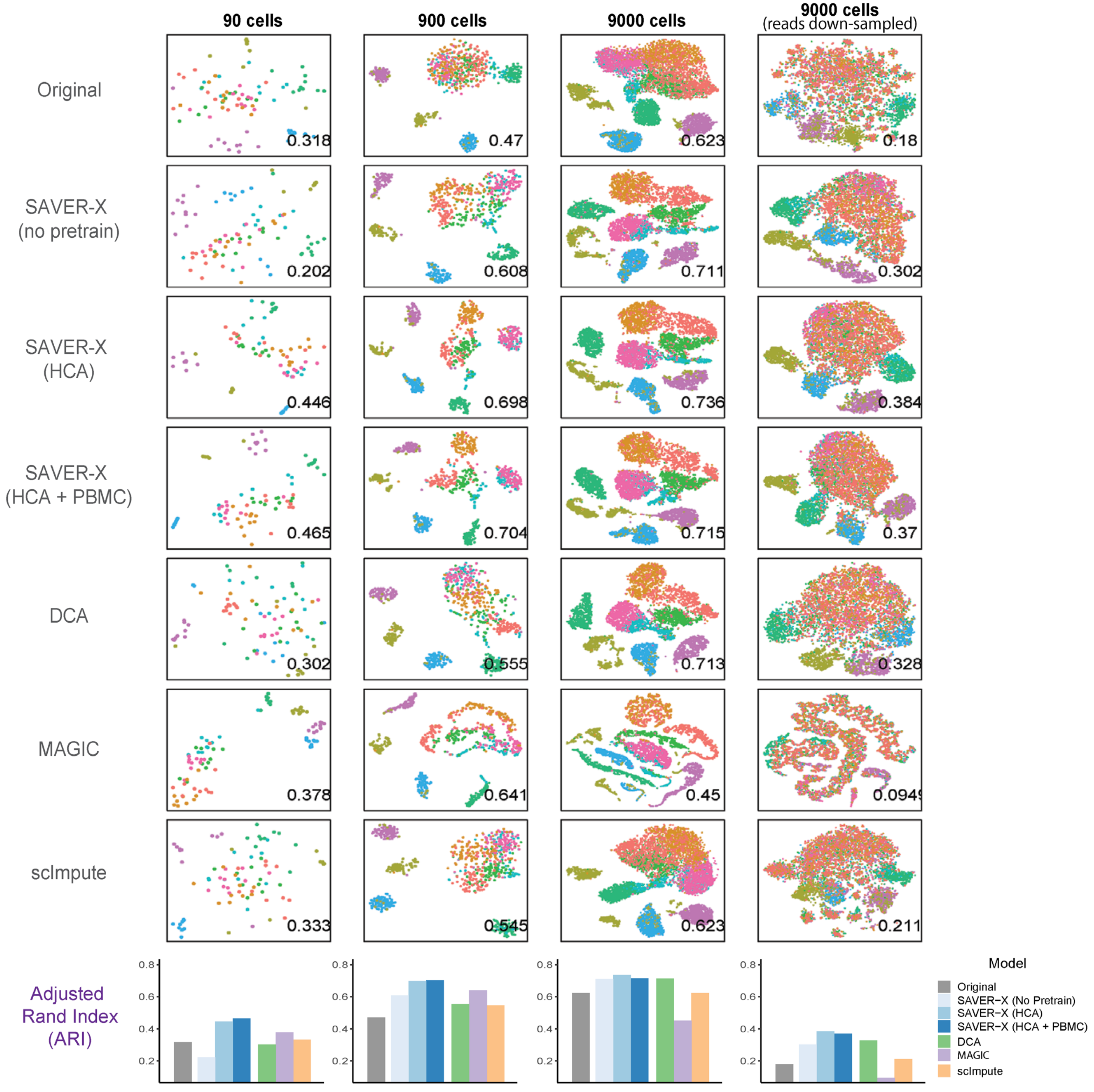


**Supplementary Figure 3**

Comparison of denoising methods under four scenarios with varying numbers of cells and the sequencing depth.

The performance of SAVER-X is benchmarked against existing denoising methods using t-SNE visualization and clustering adjusted rand index (ARI). The number at the right-bottom corner of each plot is the ARI, which is summarized also in the last row.

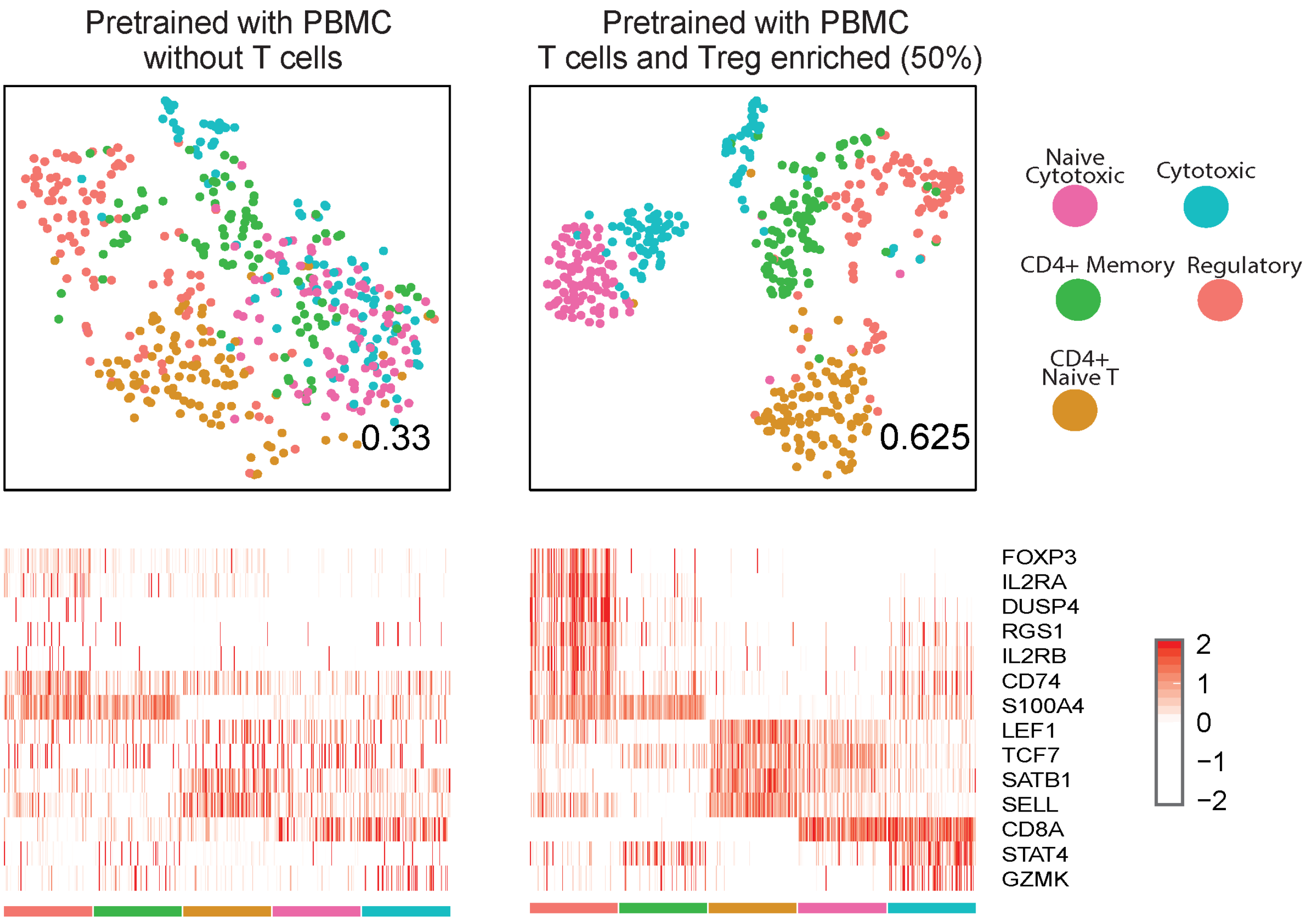


**Supplementary Figure 4**

More scenarios on PBMC T cell transfer learning.

The other two scenarios of the in-silico experiments on the 600 PBMC T cell transfer learning that is not shown in Figure 2c. More details see supplementary note 4 for more details.

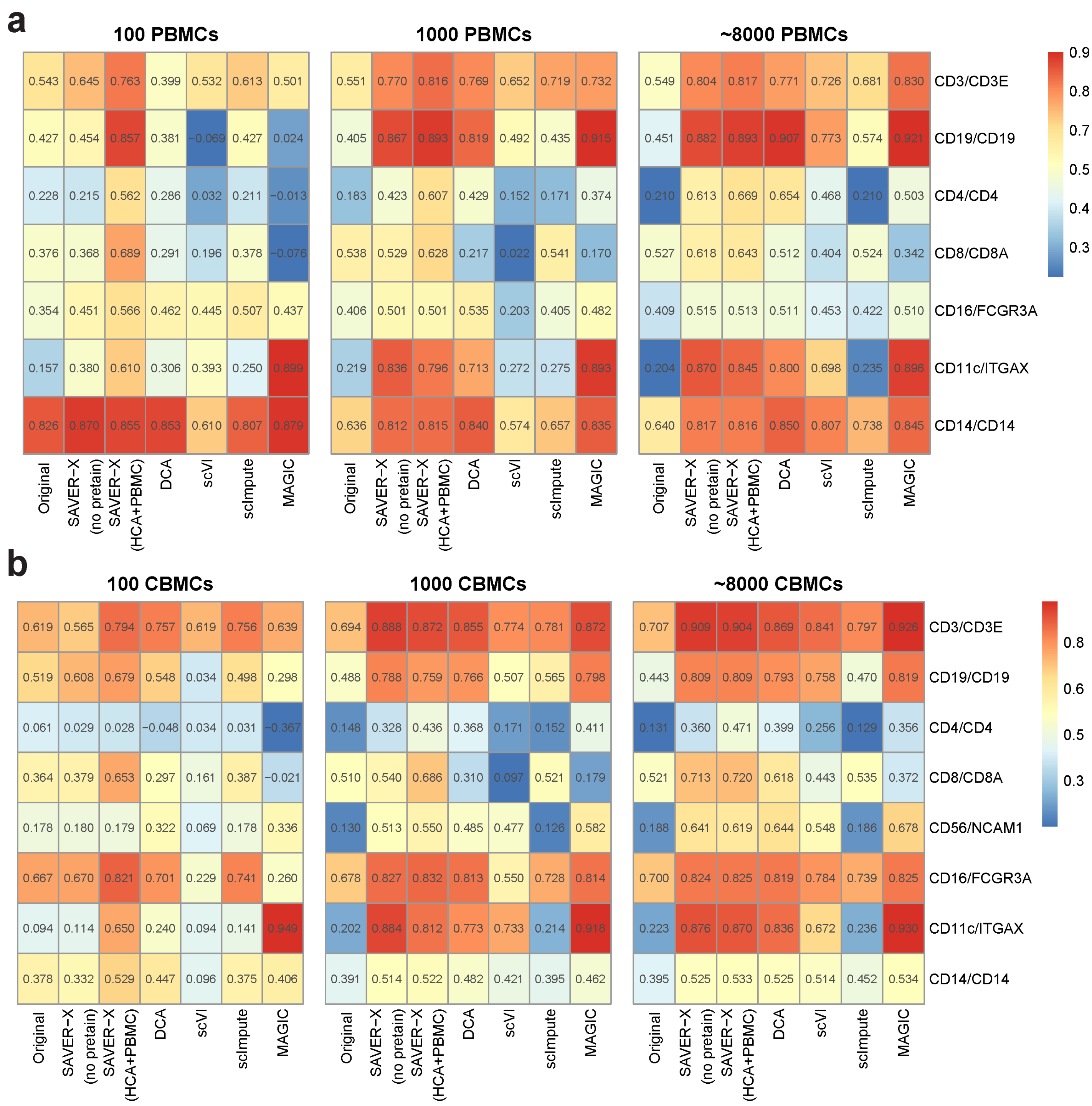


**Supplementary Figure 5**

Benchmarking scRNA-seq data denoising methods using the CITE-seq data (Stoeckius, M. et al, 2017).

Pearson correlation were calculated between proteins and corresponding mRNA levels in the CITE-seq PBMC (a) and CBMC (b) datasets. The mRNA measurements were denoised using SAVER-X and other benchmarking denoising methods (X-axis). The effect of transfer learning shows significantly when the information available from the target data is limited.

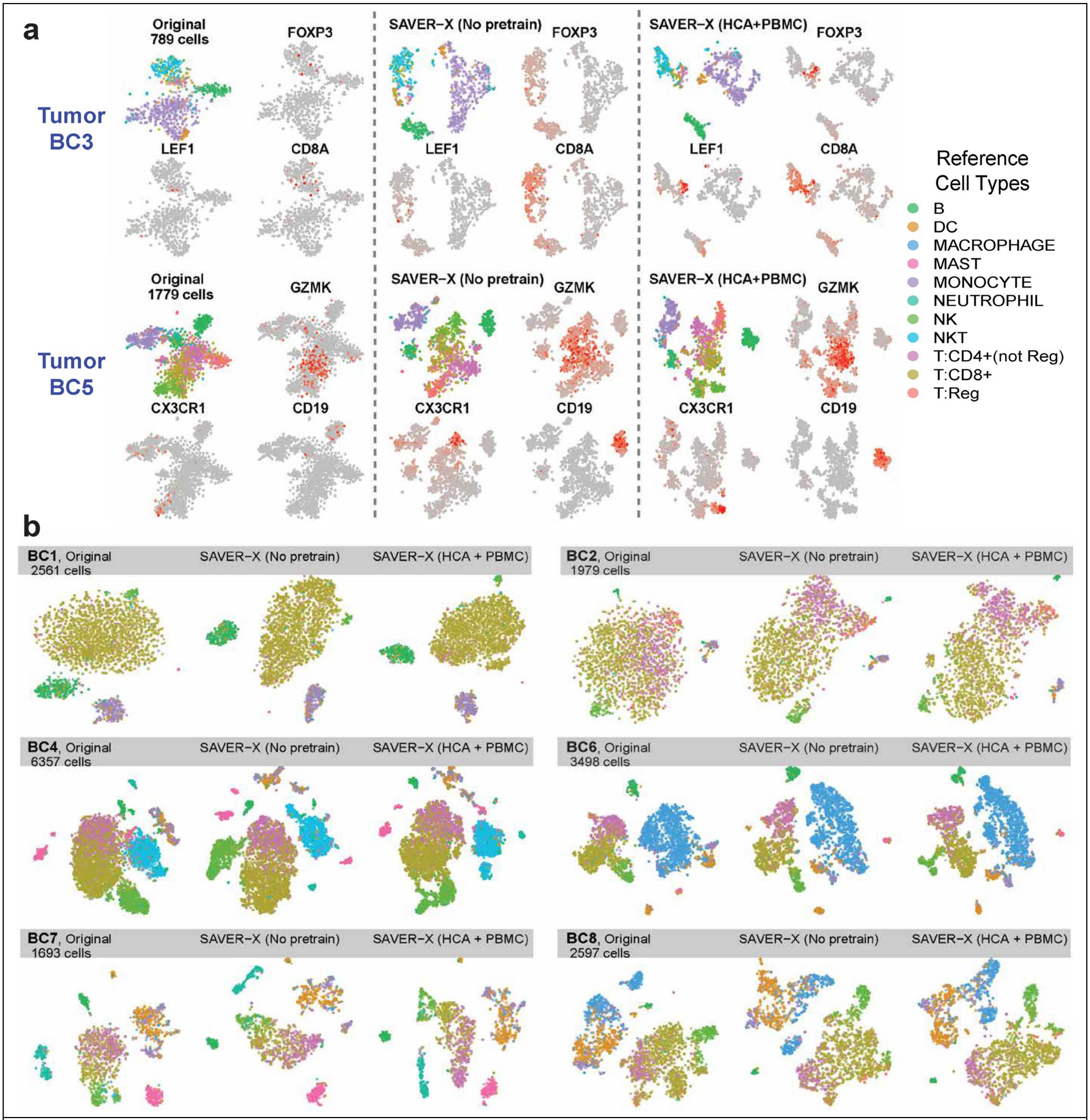


**Supplementary Figure 6**

Transfer learning from normal to breast cancer immune cells.

**a**) Infiltrating immune cells in resected breast carcinoma from two breast cancer patient tumors from Azizi et al (2018). The three panels show visualizations using original data, denoised values by SAVER-X without pretraining, and denoised values by SAVER-X pretrained on immune cells (HCA and 10X PBMC data). The t-SNE plots show separation between cell types (cell labels are obtained from the original paper). Feature plots show the expression of some known marker genes, and a darker red color represents a relatively higher expression level in some cells compared with the rest of cells. **b**) t-SNE plots for tumors of the other 6 breast cancer patients.

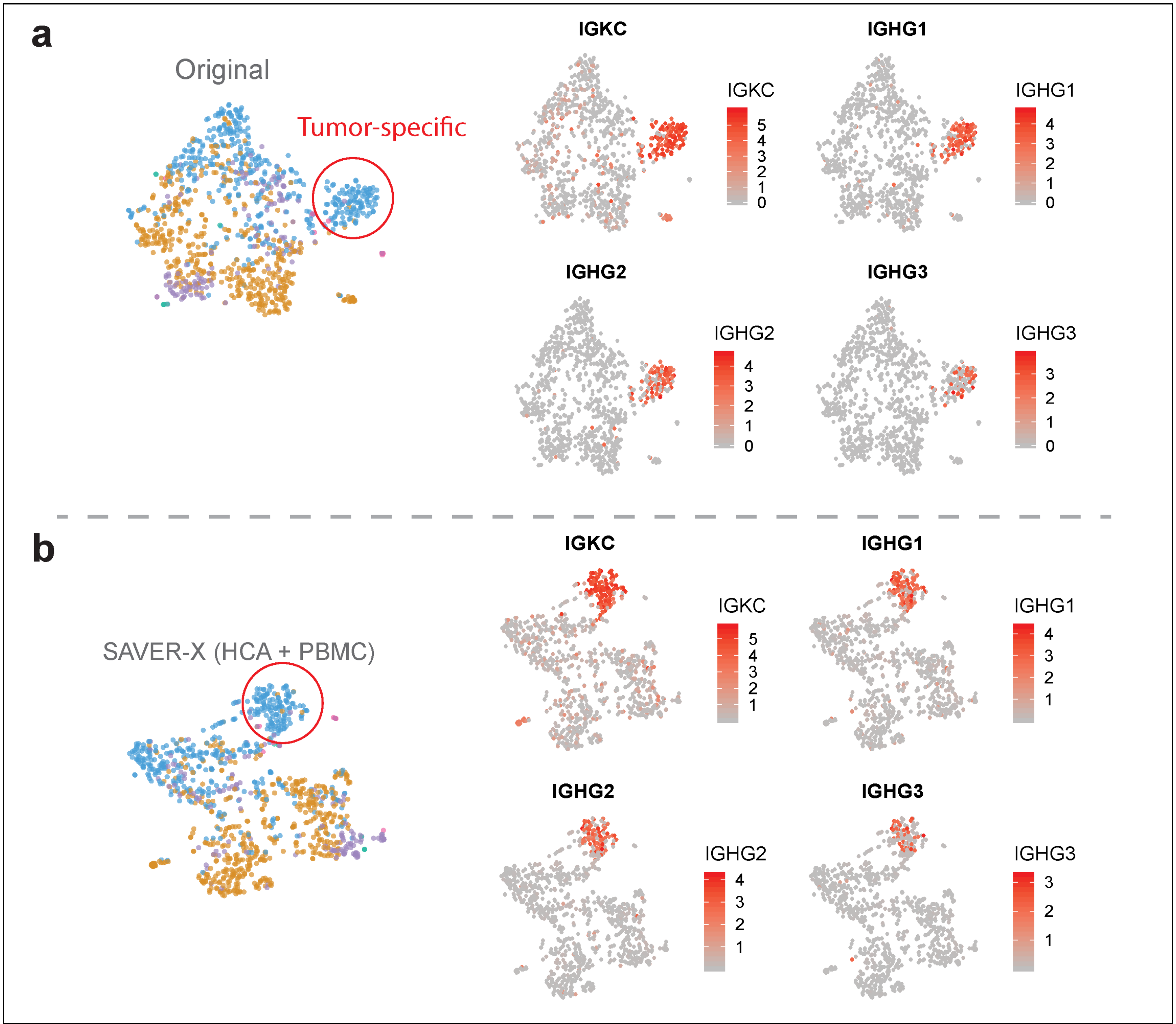


**Supplementary Figure 7**

Marker genes of the tumor-specific cell group in tumor BC8 of Azizi et al (2018).

**a**) tSNE plot and feature plots of the BC8 tumor myeloid cells (1046 cells) using the original raw data. The highly enriched expressions of immunoglobulin genes confirm that this is a tumor-related cell population. **b**) The t-SNE plot and feature plots of the BC8 tumor myeloid cells using the SAVER-X denoised data, after pretraining with the normal immune cells. A darker red color represents a relatively higher expression level in some cells compared with the rest of cells.

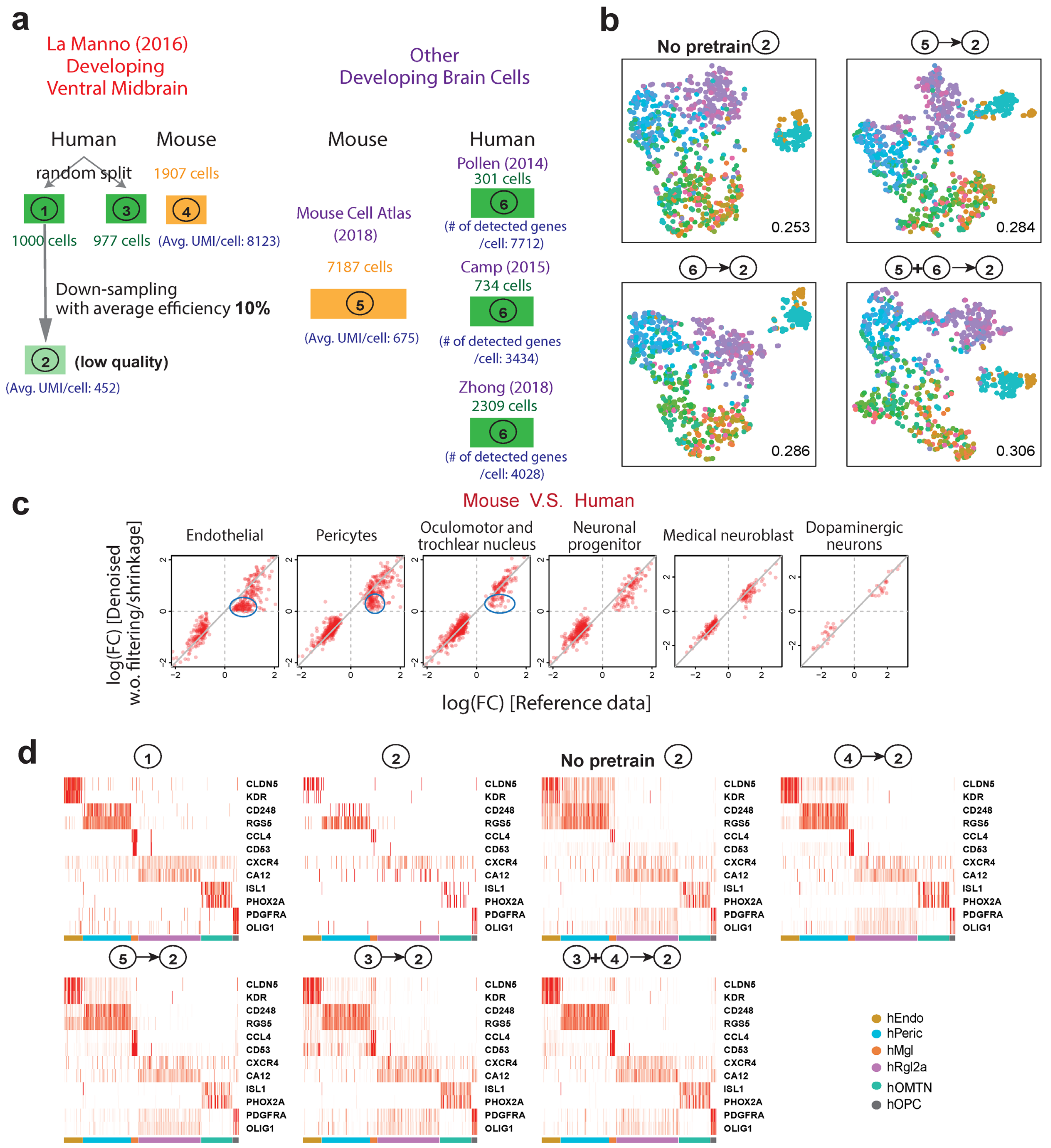


**Supplementary Figure 8**

SAVER-X analyses of the La Manno et. al. (2016) data.

**a**) Illustration of the complete design and data use. 1977 human cells are randomly split into two groups. The 1000 cells in group 1 are further down sampled. We consider pretraining with 1907 mouse cells in the same paper, 977 original human cells, 7187 mouse cells from MCA and a total of 3344 non-UMI human developmental brain human cells. **b**) t-SNE plots of the 1000 down-sampled cells for other denoising models not shown in Figure 3b. Cell labels are the computed labels from the original paper. The numbers at the right corner are the ARI for each plot. Cell types are colored the same as in Figure 3b. c) Log fold changes between human and mouse data of cell-type-specific differentially expressed genes. X axis uses the original human data and Y axis uses the denoised down-sampled human data which is denoised using SAVER-X pretrained with the paired mouse cells from La Manno et. al. (2016), but without gene filtering or empirical Bayes shrinkage. Each dot is a differentially expressed gene between human and mouse in that cell type. Genes that have notable bias are highlighted with blue circles. d) the heatmaps of the denoised gene expressions for a set of known marker genes for 5 human brain cell types.

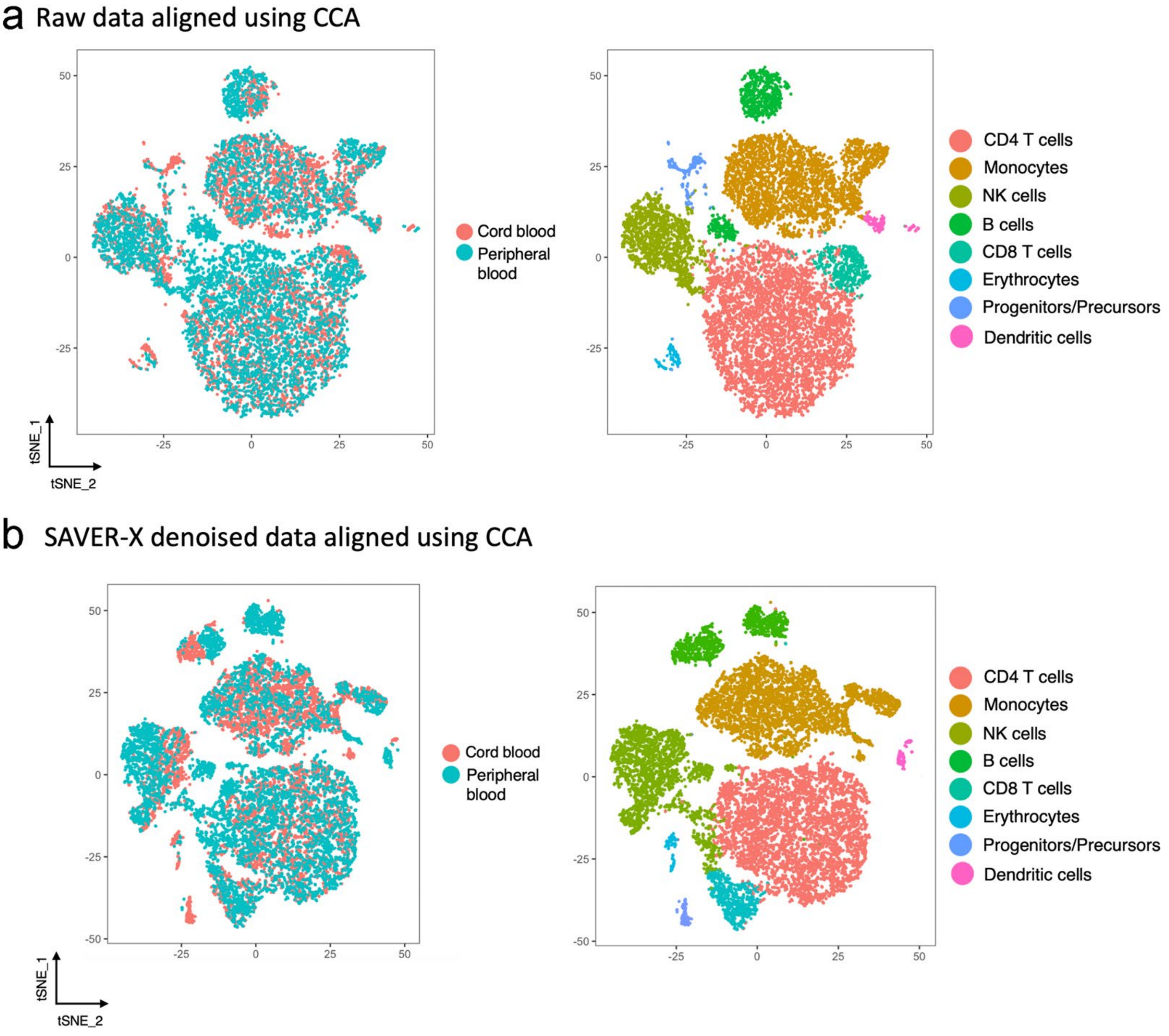


**Supplementary Figure 9**

Data alignment pre- and post-denoising yields consistent clustering results.

Panels (**a**) and (**b**) show the results of data alignment using the raw and denoised results, respectively, when we align the CBMC and PBMC scRNAs-seq datasets of ~8000 cells each. The panels on the left show the results of data alignment using 20 canonical components in SeuratCCA v2, whereas the panels on the right demonstrate the cell type identities of the clusters identified after alignment. See supplementary note 6 for more details.
